# Resolution of the D4Z4 repeat responsible for facioscapulohumeral muscular dystrophy with HiFi sequencing

**DOI:** 10.64898/2026.04.10.717730

**Authors:** Xiao Chen, Richard J. L. F. Lemmers, Zev Kronenberg, Joseph M. Devaney, Jessica Noya, April S. Berlyoung, Shamila Yusuff, Solomon Lynch, Keith Nykamp, Amanda S. Lindy, Egor Dolzhenko, Silvère M. van der Maarel, Michael A. Eberle

## Abstract

The D4Z4 macrosatellite repeat encompasses some of the most difficult-to-resolve disease-related variations in the human genome. D4Z4 has a repeat unit of 3.3 kb (encoding the *DUX4* gene) that is present in up to 100 copies on two chromosomes (4 and 10), while *DUX4* can only be expressed in somatic cells from the permissive A haplotype that usually occurs on chromosome 4. Facioscapulohumeral muscular dystrophy (FSHD) is caused by chromatin relaxation and ectopic expression of *DUX4* in skeletal muscle, mediated by contraction of D4Z4 to 1-10 copies (FSHD1, 95% of FSHD cases) or mutations in chromatin factor genes such as *SMCHD1* (FSHD2, 5% of FSHD cases). Due to its large size, disease specific haplotypes and sequence homology between chromosomes, D4Z4 is challenging to resolve by current sequencing technologies. We report a computational tool, Kivvi, to genotype D4Z4 using PacBio whole-genome long-read sequence data. Kivvi detects all D4Z4 alleles in a sample, reporting the repeat size, chromosome (4 vs. 10), distal haplotype (A vs. non-permissive haplotypes) and the methylation level of each allele. We validated Kivvi against gold standard assays for FSHD diagnostics, detecting 100% of contracted alleles and correctly classifying 90% of noncontracted alleles. We showed differential methylation signals between FSHD1 and candidate FSHD2 samples. We profiled D4Z4 across 601 individuals from five ancestral populations, revealing extensive genetic diversity. We identified common haplotypes of D4Z4 alleles and characterized hybrid repeat units, hybrid repeat arrays, and translocation alleles. Combined with HiFi long reads, Kivvi enables the consolidation of multiple FSHD assays into a single workflow and facilitates the discovery of novel genetic modifiers of FSHD through population-scale studies.

## Introduction

Facioscapulohumeral muscular dystrophy (FSHD) is one of the most prevalent inherited muscular dystrophies in adults^1,2^. In most cases, it is an autosomal dominant disease characterized by progressive weakness and atrophy of the facial, shoulder and upper-arm muscles^3,4^. FSHD exhibits a wide inter and intrafamilial clinical variability: some FSHD carriers remain non-penetrant throughout their life while others experience rapid disease progression leading to wheelchair dependency^5,6^.

FSHD is caused by ectopic expression of the *DUX4* retrogene in skeletal muscle, leading to apoptosis^7,8^. *DUX4* is a transcription factor normally silenced in somatic tissues after a brief pulse of expression during the embryonic cleavage stage^9^. A copy of the *DUX4* open reading frame is located in each unit of D4Z4, which is a variable number tandem repeat (VNTR) with a GC-rich repeat unit (RU) of 3.3 kb (Figure 1A) in the subtelomeric region of chromosome 4q35. D4Z4 occurs up to 100 copies per allele across individuals^10,11^ and normally shows high CpG methylation that correlates with the repeat size^12^, and a repressive chromatin structure^13^.

**Figure 1.**
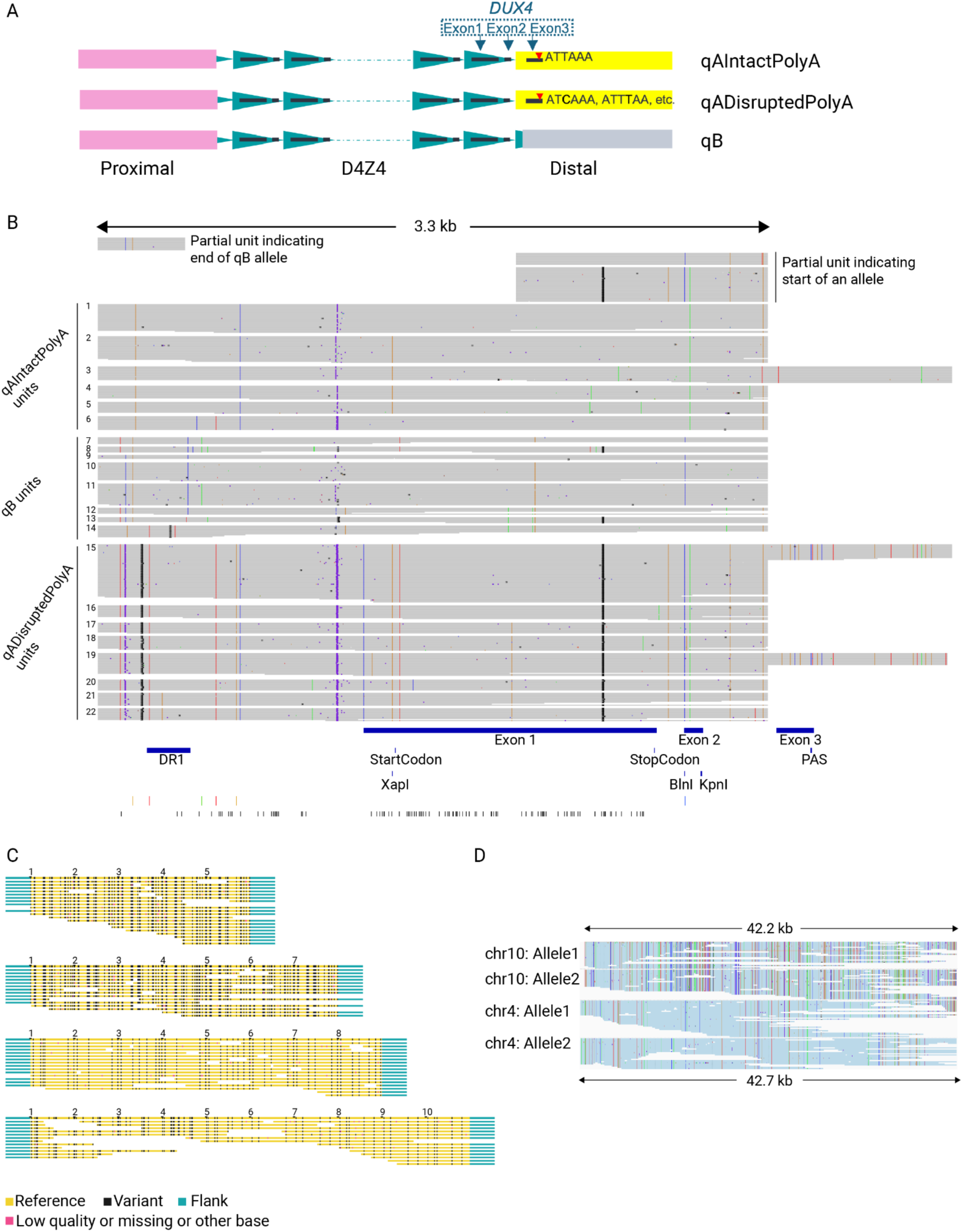
Overview of the D4Z4 locus and Kivvi. **A.** D4Z4 is a VNTR with a repeat unit of 3.3kb and is present on both chromosome 4 and chromosome 10. Each repeat unit encodes the first and second exon of the *DUX4* gene. Proximal to D4Z4, chromosome 4 and chromosome 10 share sequence homology for a ∼42 kb region. The third exon of *DUX4* is located in the sequence distal to D4Z4 and contains the somatic polyadenylation signal (PAS), denoted by the red triangle. D4Z4 alleles can be classified into three types based on the distal region: qA with an intact polyA signal (qAIntactPolyA), qA with a disrupted polyA signal (qADisruptedPolyA) and qB, where the distal sequence is different from qA and thus completely lacks the third exon and polyA signal. **B-D.** The methods used in Kivvi are shown using HG02071 as an example, which has all three types of D4Z4 alleles, each with a relatively short repeat array size. **B.** HiFi reads are realigned against one D4Z4 unit, plus a short region distal to D4Z4, including the PAS. Note that the single repeat unit used as reference here differs from the conventional definition of a D4Z4 repeat unit, which spans the region between two KpnI restriction sites. Read segments are classified into unique repeat units based on sequence variants, and these unique repeat units are labeled with distinct indices (1-22 in this sample). All alleles start with a partial unit, indicated by the shorter read segments at the top right. All qB alleles end with a partial unit indicated by the shorter read segments at the top left. All qA alleles (both qAIntactPolyA and qADisruptedPolyA) continue to the distal sequence containing the third exon. Labeled at the bottom are the exons, start and stop codons, PAS and the DR1 region (regulatory region conventionally used for methylation analysis), as well as widely used restriction sites (XapI, BlnI and KpnI). Six variant sites used for classifying the repeat unit types are labeled in colors indicating nucleotide changes (T: red, A: green, G: orange and C: blue) (See Methods). CpG sites used for methylation profiling in Kivvi are labeled in black. **C.** Repeat units are assembled into alleles based on links indicated by HiFi reads. In this image generated by Kivvi, each panel shows an allele at the top followed by its supporting reads. The boundaries of each repeat unit are marked on each allele. Informative sites are plotted with different colors indicating reference (yellow), variant (black), missing or disagreeing information (pink) and flanking sequence (teal). In this sample (HG02071), four alleles are assembled: LeftFlank-6-1-4-1-5-2-2-3-RightFlank (8RU, qAIntactPolyA), LeftFlank-14-12-13-8-7-9-10-11-10-11-RightFlankB (10RU, qB), LeftFlank-15-15-16-15-15-RightFlank (5RU, qADisruptedPolyA) and LeftFlank-22-20-19-18-21-17-19-RightFlank (7RU, qADisruptedPolyA). **D.** Kivvi uses Paraphase^34^ to phase the haplotypes in the upstream region proximal to D4Z4. WGS reads are realigned against chromosome 4 and grouped into four haplotypes, two from chromosome 10 and two from chromosome 4. The 5’ (left) ends of the haplotypes reach the end of the chromosome 4-10 homology region, where chromosome 10 haplotypes are softclipped and hence are shorter than chromosome 4 haplotypes.

FSHD patients, however, show reduced methylation and loss of repressive histone modifications, leading to partial chromatin relaxation and sporadic *DUX4* expression in skeletal muscle^14^. This chromatin relaxation can be caused by two different genetic events. First, FSHD1 (>95% of FSHD cases) is caused by a contraction of the D4Z4 array to 1-10 units^15^. Second, FSHD2 (<5% of FSHD cases) is caused by pathogenic variants in chromatin factors that establish and/or maintain the D4Z4 chromatin structure, most often *SMCHD1*^16^ and rarely *DNMT3B*^17^ or *LRIF1*^18^. The D4Z4 repeat size is known to be a major factor affecting the disease severity^19^: For FSHD1, carriers of alleles with 1-6 units are moderately to severely affected, while carriers of alleles with 7-10 units have more clinical variability and more cases of non-penetrance^20,21^.

D4Z4 is also present in the subtelomeric region of chromosome 10q26, but repeat contractions on 10q are usually not associated with FSHD^15,22^. Even on chromosome 4, only one of the two major subtelomeric variants, 4qA but not 4qB, is associated with FSHD^10,23^. This is because only D4Z4 arrays on 4qA can express *DUX4* in the skeletal muscle. The *DUX4* locus within the D4Z4 repeat unit lacks a polyadenylation signal (PAS). 4qA alleles have, immediately distal to the D4Z4 array, a somatic *DUX4* polyadenylation signal^15^, which is disrupted on 10q alleles (10qA) and missing on 4qB alleles (Figure 1A). Hence the 4qA allele is referred to as the permissive FSHD allele. The repeat size on the non-permissive alleles 4qB and 10qA can vary between 1-100 units without pathologic consequences.

D4Z4 array size determination is conventionally detected by restriction fragment digestion, linear or pulsed-field gel electrophoresis followed by Southern blotting^24^. This testing workflow is low-throughput and labor-intensive, and can sometimes give ambiguous results due to homology between chromosomes 4qA, 4qB and 10, as well as complex rearrangements in the D4Z4 locus. Optical genome mapping^25–27^ has recently been applied to replace Southern blot for sizing D4Z4 arrays. However, both of these technologies have to be combined with other assays such as bisulfite sequencing to determine the methylation level of D4Z4 arrays^28,29^, as well as sequencing of FSHD2-related genes for pathogenic variants.

Advances in long-read sequencing technologies provide an opportunity to potentially consolidate the multiple time-consuming assays for FSHD into a single workflow. Though, due to the complexity of the D4Z4 locus, de novo genome assemblies of long read data can be error prone in this region, and specialized bioinformatics strategies are needed. Previous studies have used Oxford Nanopore sequencing to genotype D4Z4 in either CRISPR/Cas9-targeted sequencing data^30^ or whole genome sequencing data^31–33^. However, these studies rely on examination of individual reads that fully contain the D4Z4 repeat, which are often longer than the read length. A comprehensive tool reporting the full set of information for D4Z4, including size, chromosome, presence of polyA signal and methylation, is desired.

Here we report a computational tool, Kivvi, to resolve D4Z4 using PacBio HiFi whole-genome sequence (WGS) data. Kivvi reports the size, sequence, originating chromosome (4 vs. 10), distal haplotype (qA vs. qB, and status of the polyA signal) and methylation level of individual D4Z4 alleles in a sample. We demonstrated differential methylation signals for distinguishing between FSHD1 and FSHD2 individuals. We then used Kivvi to profile D4Z4 alleles in 601 individuals across five ancestral populations.

Population-wide analysis of complete D4Z4 sequences allowed us to characterize common haplotypes of D4Z4, profile their sequence variations and identify hybrid repeat units, hybrid repeat arrays and translocation events.

## Results

### Overview of Kivvi

Kivvi extracts all reads overlapping D4Z4 on either chromosome 4 or chromosome 10 and realigns them to a single D4Z4 repeat unit (Figure 1B). After realignment, Kivvi identifies unique copies of the D4Z4 repeat unit and assembles them into alleles (Figure 1B and 1C). To distinguish chromosome 4 alleles from chromosome 10 alleles, Kivvi uses Paraphase^34^ to phase a 42kb genomic region that is homologous between chromosome 4 and chromosome 10 upstream of D4Z4 (Figure 1D). Kivvi reports the following information for all D4Z4 alleles of a sample: 1) distal allele type (qAIntactPolyA, qB or qADisruptedPolyA), 2) originating chromosome, 3) allele size (Kivvi reports an exact value for fully assembled alleles and a lower bound of the array size for partially assembled alleles) and 4) summary methylation level. Details of the Kivvi method are described in the Methods section.

### Validation of D4Z4 calls against orthogonal assays

Pulsed-field gel electrophoresis (PFGE)-Southern blot results are available for 27 control samples from the 1000 Genomes Project (1kGP)^35^ (Table S1) and optical genome mapping (OGM) results are available for 10 FSHD1 patient samples (Table S2). In this paper, we refer to these two datasets as the 1kGP dataset and the FSHD1 dataset, respectively. Kivvi results are summarized separately for these two datasets due to the difference in the HiFi data quality (Figure S1), assessed by two key metrics for assembly: read length and read depth in the D4Z4 region. The 1kGP samples have a median read length of 15-20kb and a depth of 10-15X per D4Z4 allele, while the FSHD1 samples have a median read length of 13-15kb and a per-allele depth of 5-10X (Figure S1). Since this region is difficult to assemble, shorter read length and lower sequencing depth will make samples more prone to performance degradation.

Across both datasets, Kivvi correctly classified 147/148 (99.3%) of alleles into qA with intact polyA (hereafter referred to qAIntactPolyA), qB or qA with disrupted polyA (hereafter referred to qADisruptedPolyA). One expected qADisruptedPolyA allele was not reported by Kivvi due to the high sequence similarity between two qADisruptedPolyA alleles in the same sample (two alleles merged into one at the distal end).

Next, we assessed the accuracy of Kivvi reported allele sizes. Kivvi reports an exact size for fully assembled alleles and a lower bound of the repeat array size for partially assembled alleles (see Methods). Across both datasets, all alleles fully assembled by Kivvi were consistent with the expected size allowing a difference of up to three units (Table S1 and S2). To broaden our assessments for partially assembled alleles, where a reported lower bound higher than 10 is meaningful for FSHD testing, we consider two scenarios in Table 1. First, a truth allele of 1-10 repeat units (RU), hereafter referred to as 1-10RU, is assigned “fully assembled with correct size” if it is fully assembled with a matching size by Kivvi. It is considered inconclusive (for testing purpose) if it’s not fully assembled. Second, a truth allele longer than 10 RU, hereafter referred to as >10RU, is assigned “fully assembled with correct size” if it is fully assembled with a matching size by Kivvi. A >10RU truth allele partially assembled by Kivvi is assigned “partially assembled and correctly classified” if the reported lower bound is longer than 10, and is otherwise considered inconclusive.

**Table 1.** Validation of D4Z4 calls against PFGE-Southern blot in 27 1kGP samples and against Optical Genome Mapping in 10 FSHD1 samples. Concordance is calculated as (fully assembled with correct size + partially assembled and correctly classified)/total alleles.

|  | Allele type | Allele size (RU) | Total alleles | Fully assembled with correct size | Partially assembled and correctly classified | Inconclusive | Concordance |
| --- | --- | --- | --- | --- | --- | --- | --- |
| 1kGP dataset | qAIntactPolyA | 1-10 | 1 | 1 | 0 | 0 | 1 |
|  |  | >10 | 30 | 6 | 21 | 3 | 0.9 |
|  |  | total | 31 | 7 | 21 | 3 | 0.903 |
|  | qB | 1-10 | 6 | 6 | 0 | 0 | 1 |
|  |  | >10 | 20 | 14 | 6 | 0 | 1 |
|  |  | total | 26 | 20 | 6 | 0 | 1 |
|  | qADisruptedPolyA | 1-10 | 6 | 4 | 0 | 2 | 0.667 |
|  |  | >10 | 45 | 9 | 29 | 7 | 0.844 |
|  |  | total | 51 | 13 | 29 | 9 | 0.824 |
| FSHD1 dataset | qAIntactPolyA | 1-10 | 10 | 10 | 0 | 0 | 1 |
|  |  | >10 | 4 | 0 | 2 | 2 | 0.5 |
|  |  | total | 14 | 10 | 2 | 2 | 0.857 |
|  | qB | 1-10 | 0 | 0 | 0 | 0 | N/A |
|  |  | >10 | 6 | 0 | 4 | 2 | 0.667 |
|  |  | total | 6 | 0 | 4 | 2 | 0.667 |
|  | qADisruptedPolyA | 1-10 | 4 | 1 | 0 | 3 | 0.25 |
|  |  | >10 | 16 | 2 | 10 | 4 | 0.75 |
|  |  | total | 20 | 3 | 10 | 7 | 0.65 |

In the 1kGP dataset (Table 1), there is one 1-10RU qAIntactPolyA allele and this was fully assembled with the correct size. 27 (90%) out of 30 >10RU qAIntactPolyA alleles were either fully assembled with the correct size or partially assembled and correctly classified. The three inconclusive alleles were reported as >9, >=8 and >6 RU (Table S1). All qB alleles were either fully assembled with the correct size or partially assembled and correctly classified. For qADisruptedPolyA alleles, 4 (66.7%) out of 6 1-10RU alleles were fully assembled with the correct size and 38 (84.4%) out of 45 >10RU alleles were either fully assembled with the correct size or partially assembled and correctly classified. qADisruptedPolyA alleles are expected to be more difficult to assemble since qADisruptedPolyA units are highly similar to each other (Figure S2). As qADisruptedPolyA alleles are not pathogenic when contracted, a lower performance for qADisruptedPolyA alleles does not affect FSHD testing.

The FSHD1 dataset (Table 1) showed lower performance, especially for longer (>10RU) alleles, than the 1kGP dataset due to the lower data quality. For qAIntactPolyA alleles, 10 (100%) out of 10 1-10RU alleles were fully assembled with the correct size, and 2 (50%) out of 4 >10RU alleles were partially assembled and correctly classified. Both inconclusive alleles were reported as >=9 RU (Table S2). All 6 qB alleles were expected to be >10RU and 4 (66.7%) of them were partially assembled and correctly classified. For qADisruptedPolyA alleles, 1 (25%) out of 4 1-10RU alleles were fully assembled with the correct size and 12 (75%) out of 16 >10RU alleles were either fully assembled with the correct size or partially assembled and correctly classified.

Across both datasets, Kivvi successfully reported 11 out of 11 contracted (1-10RU) qAIntactPolyA alleles, demonstrating its ability to detect pathogenic alleles in the FSHD1 size range.

### D4Z4 sizes and methylation levels in FSHD patient samples and the general population

We genotyped D4Z4 alleles with Kivvi in 601 unrelated population samples from five ancestral populations (291 Europeans, 95 Africans, 114 Admixed Americans, 54 East Asians and 47 South Asians). Among assembled chromosome 4 alleles, 40.1% are qAIntactPolyA, 57.6% are qB and 2.3% are qADisruptedPolyA (Figure S3). Among assembled chromosome 10 alleles, 0.51% are qAIntactPolyA, 3.6% are qB and 95.9% are qADisruptedPolyA (Figure S3). Hence, chromosome 4 alleles are rarely qADisruptedPolyA and chromosome 10 alleles are rarely qAIntactPolyA. Note that these frequencies might deviate from the actual frequencies in the population as the level of difficulty in assembling these three types of alleles is different (Figure S2).

In addition to the 10 FSHD1 patient samples described in the Validation section above, we also had access to 6 samples with pathogenic variants in *SMCHD1* (rows 1 to 6 in Table 2). We compared the distribution of D4Z4 sizes vs. methylation levels in the population samples, the FSHD1 samples and the samples with *SMCHD1* pathogenic variants. Note that all cell lines (280 population samples) were excluded in this methylation analysis as the methylation status is known to be variable and can be unreliable in lymphoblastoid cell lines.

**Table 2.** Kivvi results in samples with pathogenic variants in *SMCHD1*. Sample GeneDx001 (last row) is from our FSHD1 dataset and could be a FSHD1+2 patient based on the 9RU qAIntactPolyA allele, the *SMCHD1* pathogenic variant and the low overall D4Z4 methylation.

| Sample ID | *SMCHD1* pathogenic variant | D4Z4 median methylation | Number of qAIntactPolyA alleles | qAIntactPolyA allele size |
| --- | --- | --- | --- | --- |
| GeneDx011 | c.1910_1922delATGATGG AGAAGT p.H637fs HET (chr18:2705760, GRCh38) | 0.29 | 1 | >=33 |
| GeneDx012 | c.181dup p.C61Lfs HET (chr18:2656255, GRCh38) | 0.529 | 1 | >=30 |
| GeneDx013 | c.180_183delGTGT p.C61Rfs HET (chr18:2656255, GRCh38) | 0.365 | 1 | >=12 |
| GeneDx014 | c.1030 C>T p.R344* HET (chr18:2694683, GRCh38) | 0.404 | 0 | N/A |
| GeneDx015 | c.2412del p.Y804* HET (chr18:2718388, GRCh38) | 0.506 | 0 | N/A |
| GeneDx016 | c.1844 T>G p.V615G HET (chr18:2705695, GRCh38) | 0.322 | 1 | >8 |
| GeneDx001 | c.1843 G>T p.V615F HET (chr18:2705694, GRCh38) | 0.467 | 2 | 9, >=9 |

The D4Z4 repeat is generally highly methylated in unaffected individuals (Figure 2). For fully assembled alleles (Figure 2A), there is a correlation between allele size and methylation level, especially for qAIntactPolyA and qB alleles. Contracted qAIntactPolyA alleles in FSHD1 samples are distinct from other alleles with low (<0.5) methylation levels. Among the population samples, we identified one sample with a contracted (7RU), hypomethylated qAIntactPolyA D4Z4 allele (Figure 2A). This individual is not reported with FSHD, and could be a case of non-penetrance. No contracted qAIntactPolyA alleles are found in the six samples with *SMCHD1* pathogenic variants, while the reported qB and qADisruptedPolyA alleles in these samples show lower methylation levels than expected based on the size (Figure 2A).

**Figure 2.**
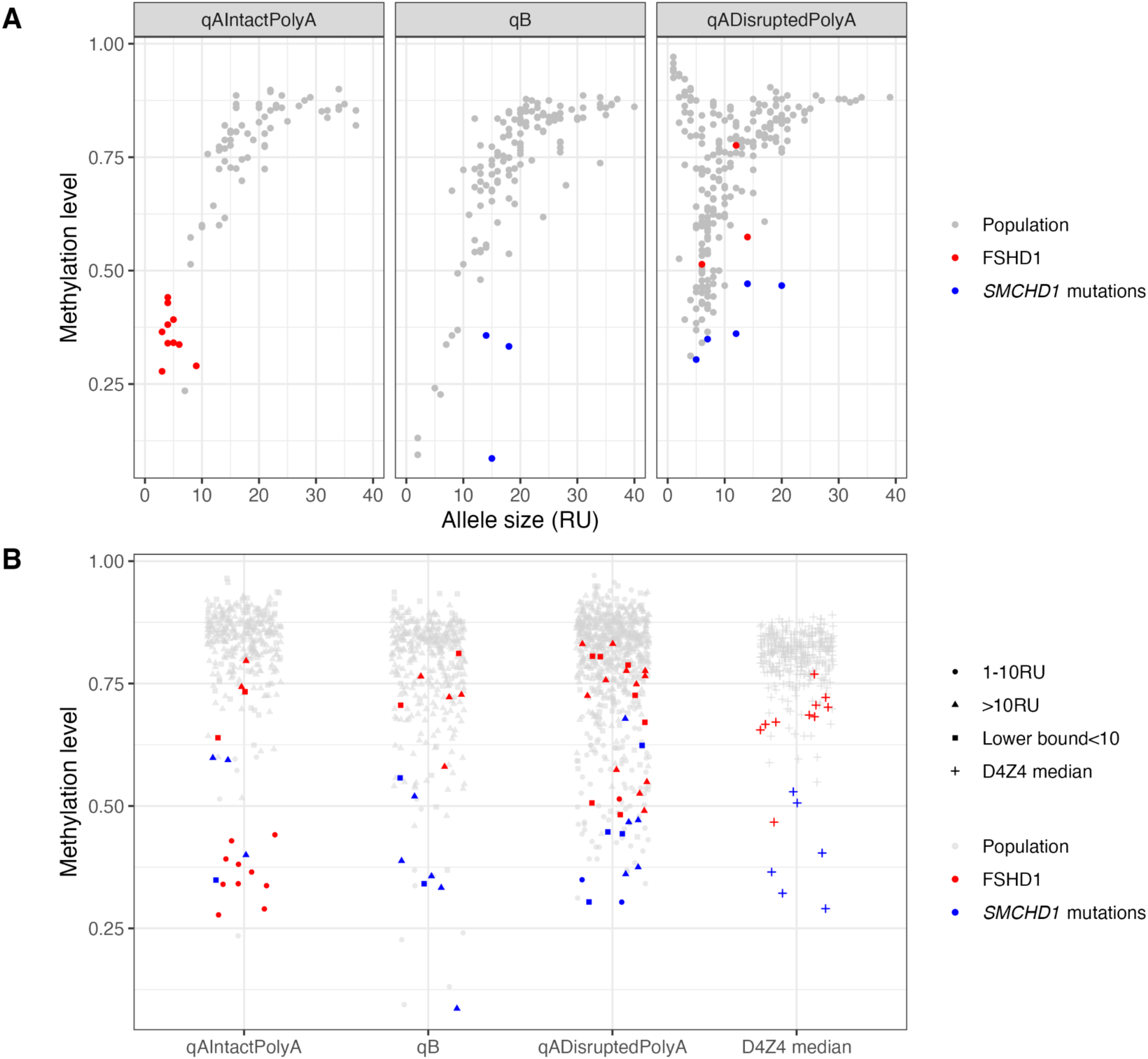
Size and methylation levels of D4Z4 alleles in population samples, FSHD1 patient samples and individuals with *SMCHD1* pathogenic variants. Distribution of allele size and methylation levels in fully assembled alleles. The methylation level of an allele is calculated as the median of all CpG sites in the last five repeat units. **B.** Methylation levels of all D4Z4 alleles, including partially assembled alleles. The data points marked as “lower bound<10” indicate a partial allele where the reported lower bound of allele size is shorter than 10 and thus cannot be classified into 1-10RU or >10RU. The last column plots the median methylation level of all D4Z4 units for each sample.

When we consider all D4Z4 alleles reported by Kivvi (Figure 2B), including partially assembled alleles, the alleles in FSHD1 samples are hypomethylated only in the contracted qAIntactPolyA alleles, while other noncontracted alleles remain highly methylated. For samples with *SMCHD1* pathogenic variants, most D4Z4 alleles are hypomethylated, regardless of the allele type and allele size. We calculated the median methylation level across all D4Z4 units as a summary metric for D4Z4 methylation of a sample (Figure 2B, fourth column). Samples with *SMCHD1* pathogenic variants show lower values for this metric, consistent with the fact that all D4Z4 alleles are affected by deficiencies in the epigenetic machinery that represses the D4Z4 chromatin structure. In contrast, FSHD1 samples have higher values for this metric as alleles beyond the

FSHD1 size range are all highly methylated. The overall D4Z4 methylation of one FSHD1 sample, however, falls within the range of samples with *SMCHD1* pathogenic variants (Figure 2B). We further checked this sample and identified a potentially pathogenic variant in *SMCHD1* (last row in Table 2), located in the same codon as one of the six samples with *SMCHD1* pathogenic variants (Figure S4). The contracted qAIntactPolyA allele in this sample is 9 RU, which falls in a size range indicating milder phenotypes for FSHD1. This individual is potentially FSHD1+2, consistent with previous studies reporting overlaps between FSHD1 and FSHD2 patients^36,37^. Indeed, approximately 10% of FSHD2 patients carry a qAIntactPolyA allele with 8-10 RU^36,37^.

Figure 3 plots the methylation levels of individual repeat units across D4Z4 alleles for the three allele types (Figure 3A-C), highlighting hypomethylation of contracted qAIntactPolyA alleles (Figure 3A). Consistent with findings from previous studies^32^, proximal units are less methylated than distal units (Figure 3D).

**Figure 3.**
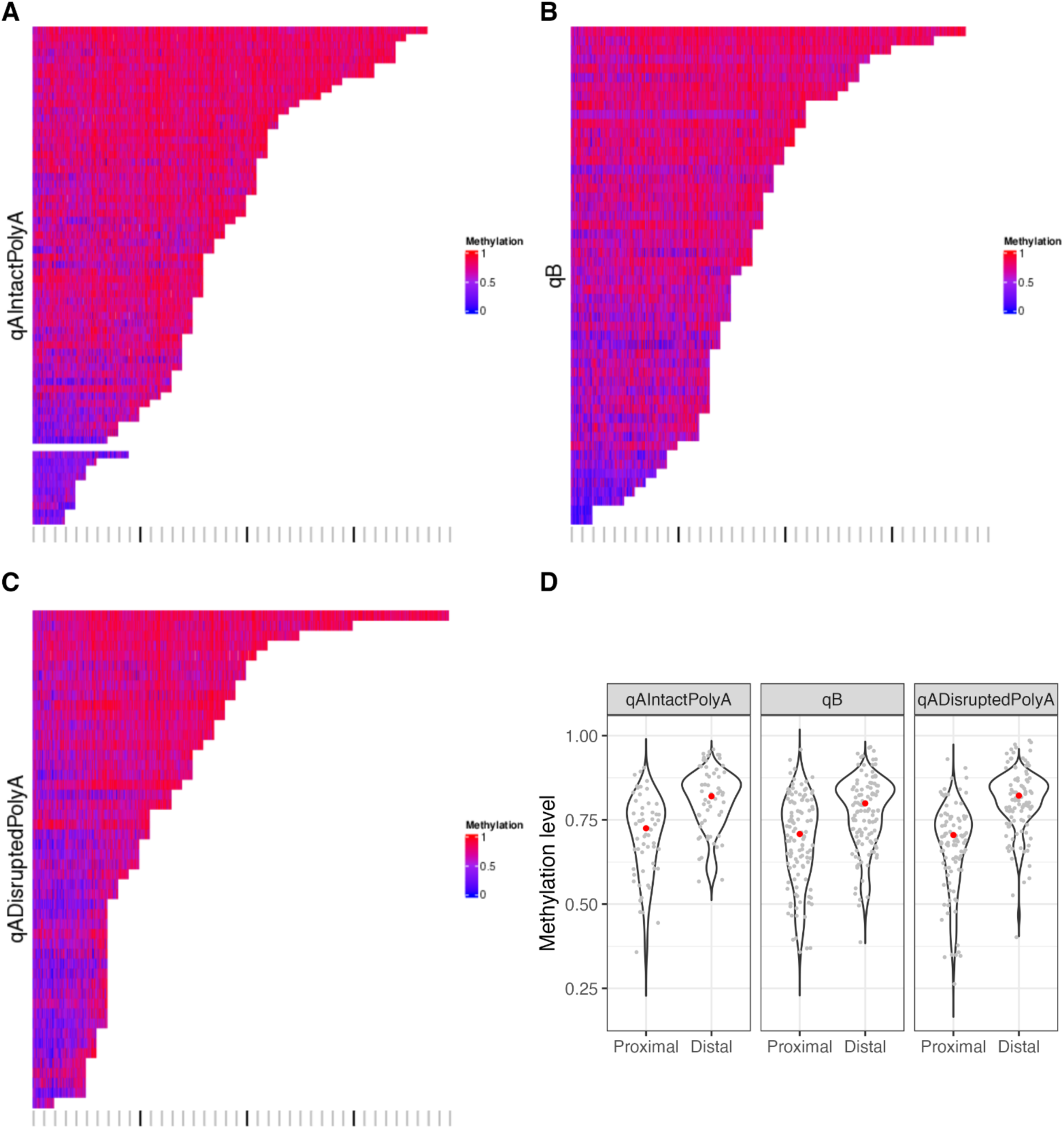
Methylation across repeat units in D4Z4 alleles. **A-C.** Methylation levels plotted across repeat units in individual D4Z4 alleles for qAIntactPolyA (A), qB (B) or qADisruptedPolyA (C) alleles. Contracted qAIntactPolyA alleles in FSHD1 patient samples are shown at the bottom of panel A. For each repeat unit, 101 CpG sites are plotted (see Methods). The gray and black bars at the bottom of each panel mark the repeat sizes, with black bars representing size 10, 20 and 30. **D.** Comparison of methylation levels at the proximal end vs. the distal end of D4Z4 arrays. For each fully assembled allele of 10 units or longer, the median methylation values of 5 most proximal units and the median of 5 most distal units are plotted.

### Sequence variations of D4Z4 alleles across populations

We analyzed 14,325 D4Z4 repeat units from 791 fully assembled D4Z4 alleles and performed principal component analysis (PCA) on these repeat units based on their sequence variants. The allele type of a D4Z4 allele is defined by its distal end (qAIntactPolyA, qB or qADisruptedPolyA), and every repeat unit on an allele is labeled the same as the allele type. PCA separated these repeat units into three clusters that roughly correspond to qAIntactPolyA, qB and qADisruptedPolyA (Figure 4A). We refer to these three clusters as qAIntactPolyA-like, qB-like and qADisruptedPolyA-like repeat units, defined by their distinct variant profiles (Table S3). Note that the definition differs between a qAIntactPolyA-like (or qB-like or qADisruptedPolyA-like) repeat unit (defined by the variants within the repeat unit) and a qAIntactPolyA (or qB or qADisruptedPolyA) repeat unit (defined by the distal end of the allele containing the repeat unit).

**Figure 4.**
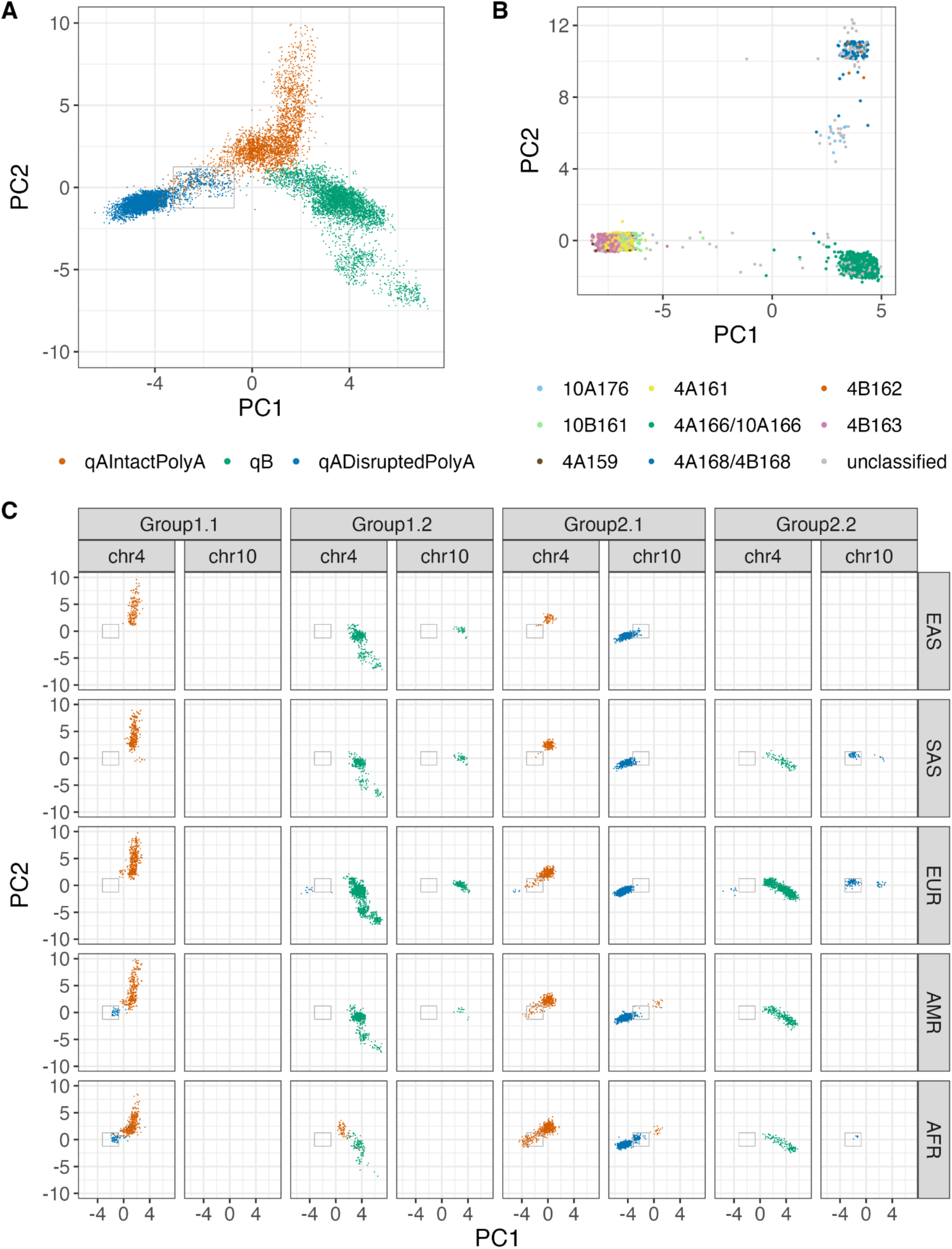
Sequence variations of D4Z4 repeat units and upstream regions across populations. **A.** Principal component analysis separates D4Z4 repeat units into three clusters. Every repeat unit on a qAIntactPolyA, qB or qADisruptedPolyA allele is labeled as qAIntactPolyA, qB or qADisruptedPolyA, respectively. The gray-bordered box denotes the qAIntactPolyA-qADisruptedPolyA hybrid zone. **B.** Principal component analysis of haplotypes from a 3640bp region directly upstream of D4Z4 identifies clusters that correspond to previously identified haplotypes. **C.** Same PCA plot as in A, showing repeat units grouped by population, chromosome and upstream haplotypes. EAS: East Asians, SAS: South Asians, EUR: Europeans, AMR: Admixed Americans, AFR: Africans.

There is a region transitioning from qADisruptedPolyA-like to qAIntactPolyA-like repeats (Figure 4A), indicating hybrid repeat units. If we define a hybrid zone (gray-bordered box in Figure 4A), 36.9% of qAIntactPolyA alleles and 13.2% of qADisruptedPolyA alleles have at least one unit that falls into the hybrid zone. Previous studies using Southern blot analysis of D4Z4 repeat arrays^35^ described hybrid D4Z4 arrays consisting of a mixture of 4q-type (qAIntactPolyA-like) and 10q-type (qADisruptedPolyA-like) units based on the sensitivity to the restriction enzymes BlnI (B) and XapI (X), which correspond to two variants, T/C at position 2889 and C/G at position 1450, respectively. In previous studies, B-X+ units were considered 4q-type units and B+X-units were considered 10q-type units. In this study, all variants within the repeat unit are included for analysis, thus capturing more sequence variations beyond the two restriction sites.

The hybrid repeat units in the hybrid zone could indicate the transition between the previously defined 4q (B-X+) and 10q-type (B+X-) units. Indeed, B-X+ units on chromosome 4 and B+X-units on chromosome 10 rarely fall into the hybrid zone (Figure S6), while the repeat units in the hybrid zone most often have nonstandard configurations of the restriction sites, such as B-X-, B+X+, or B-X+ on chromosome 10, or B+X-on chromosome 4 (Figure S6).

Going beyond sequences of the repeat units, we performed PCA on the sequences of a 3.6kb region upstream of D4Z4. As labels, we mapped our D4Z4 alleles to previously identified haplotypes based on their reported sequence signatures^35^ (Figure 4B and Figure S7). PCA identified several clusters of upstream haplotypes that correspond to previously identified haplotypes^35^, namely, a cluster of 4A166/10A166 (bottom right), a cluster of 4A168/4B168 and 4B162 (upper right), a cluster of 4B163, 4A161, 10B161 and 4A159 (bottom left) and a cluster of 10A176 (middle right). For simplicity, here we define two major groups of upstream haplotypes (separated on PC1), each with two subgroups (separated on PC2 and PC3, respectively) (See Methods and Figure S8).

We used the upstream haplotype group and chromosome information to divide the PCA plot in Figure 4A into multiple panels in Figure 4C, showing a comprehensive set of information (from upstream region to internal repeat units and to the distal polyA signal) for each D4Z4 allele in five ancestral populations. Groups 1 and 2 define two major haplotypes in the D4Z4 region. Within each group, subgroups 1 and 2 separate qA from qB. While chromosome 4 alleles are present on all four subgroups, chromosome 10 alleles are never found on Group1.1. On chromosome 4, the qAIntactPolyA units on Group1.1 and the qAIntactPolyA units on Group2.1 occupy different spaces in the PCA plot, suggesting sequence differences between these two sets of qAIntactPolyA units. The qAIntactPolyA units on Group2.1, which correspond to previously identified haplotype 4A166, are closer to qADisruptedPolyA units and touch the hybrid zone. On chromosome 10, qADisruptedPolyA alleles are mainly on Group2.1 and occasionally on Group2.2.

In African populations, more qAIntactPolyA and qADisruptedPolyA units on Group2.1 appear in the hybrid zone (Figure 4C). In European/East Asian/South Asian populations, 29.2% of qAIntactPolyA alleles have at least one unit that falls into the hybrid zone, while in African populations, 47.5% of qAIntactPolyA alleles have at least one unit that falls into the hybrid zone. In addition to hybrid repeat units, there also exist hybrid alleles. Here, a hybrid allele refers to an allele that does not conform to the standard relationship between chromosome, upstream haplotype group, internal unit types and distal allele type. African populations have more hybrid alleles: some chromosome 4-Group1.1 alleles have the qADisruptedPolyA distal end and some chromosome 10-Group2.1 alleles have the qAIntactPolyA distal end (Figure 4C).

In addition to hybrids involving qAIntactPolyA and qADisruptedPolyA alleles, there also exist hybrid alleles involving qB-like units. We identified two chromosome 4-qADisruptedPolyA hybrid alleles, which start with qB-like units in the proximal end and transition to qADisruptedPolyA-like units in the distal end (Figure S9), indicating translocation events, where the transition point corresponds to the translocation breakpoint. Another type of hybrid alleles involving qB-like units are the chromosome 10-Group2.2 alleles (10A176), which often transition from units in the qAIntactPolyA-qADisruptedPolyA hybrid zone in the proximal end to qB-like units in the distal end (Figure S10), while the distal polyA signal is qADisruptedPolyA.

## Discussion

In this paper, we use Kivvi to study the D4Z4 repeat, one of the most difficult, clinically relevant regions of the human genome. For each sample, Kivvi calls the size (exact size or lower bound), originating chromosome, distal allele type (qAIntactPolyA, qB or qADisruptedPolyA) and methylation level for all D4Z4 alleles, providing a comprehensive summary of the information required for FSHD testing.

Comparing against PFGE-Southern blot and OGM results, Kivvi correctly identified 11 out of 11 contracted (FSHD1-sized) qAIntactPolyA alleles. In the 1kGP dataset with higher data quality, Kivvi reported informative allele sizes in 90% of noncontracted qAIntactPolyA alleles and 100% of qB alleles. Even in alleles where the Kivvi reported sizes are marked as inconclusive (Table 1), the lower bounds reported (>9, >=8 and >6) were high enough to exclude alleles of 1-6RU, which represent 61.2% of European FSHD cases and 94.3% of Asian FSHD cases based on published frequencies^38^.

Performance for qADisruptedPolyA alleles is lower, but qADisruptedPolyA alleles are not associated with FSHD and thus are not clinically informative. Our data highlight that Kivvi’s performance, particularly for assembling long (>10RU) alleles, can be influenced by data quality, most importantly read depth and read length (Table 1 and Figure S1), and would benefit from increased HiFi read length in the future.

Kivvi also profiles the methylation levels of D4Z4 alleles by leveraging the 5-methylcytosine (5mC) information in HiFi reads. Kivvi is designed to identify both FSHD1 and FSHD2 cases with different characteristics. FSHD1 cases have a 1-10RU qAIntactPolyA allele that is hypomethylated, while other non-contracted D4Z4 alleles remain highly methylated (Figure 2). In contrast, FSHD2 cases, most often caused by a pathogenic variant in *SMCHD1*, are marked by overall reduction in D4Z4 methylation, with all D4Z4 alleles hypomethylated (Figure 2). In this study, we only included seven samples with *SMCHD1* pathogenic variants (including the one candidate FSHD1+2 sample). Future studies are needed to test confirmed FSHD2 cases with Kivvi and measure Kivvi’s accuracy in calling D4Z4 alleles in the 11-20 size range commonly indicated for FSHD2.

We profiled the D4Z4 repeat sequences in 601 individuals across five ancestral populations. PCA clusters D4Z4 repeat units into three clusters that correspond to the three allele types (qAIntactPolyA, qB or qADisruptedPolyA) (Figure 4A). Pairwise distance calculation (Figure S2) shows that the qB units are the most heterogeneous while qADisruptedPolyA units are highly similar to each other, consistent with previous observations^10^. As a result, qADisruptedPolyA alleles are most difficult to assemble (Table 1). Pairwise distance between two units on the same allele is lower than that between any pair of units of the same allele type in the same population (Figure S2).

Repeat units on the same allele are often products of duplication and can interact with each other through gene conversion, and hence are more similar in sequence.

In our PCA (Figure 4C), we compared the full set of information (from upstream region to internal repeat units and to the distal polyA signal) for each D4Z4 allele in five ancestral populations. Each combination of the upstream haplotype group and the originating chromosome usually defines a unique type of D4Z4 allele. Deviations from this chromosome-upstream group-allele type relationship allowed us to identify hybrid alleles, some of which show strong signals for a translocation event, transitioning from one type of units to another type of units (Figure S9). In addition to hybrid alleles, the qAIntactPolyA-qADisruptedPolyA hybrid zone (Figure 4A) contains repeat units with mixed sequence signatures, which could suggest some interactions between qAIntactPolyA and qADisruptedPolyA alleles or are remnants of the evolution toward the different D4Z4 haplotypes. More hybrid units and hybrid alleles are found in African populations (Figure 4C), as was shown in previous studies^35^, consistent with greater genetic diversity in individuals with African ancestry. These complete D4Z4 sequences provide new opportunities for population-scale analyses to understand the evolution of this dynamic region, as well as to help refine disease mechanisms and unveil new genetic modifiers of FSHD.

D4Z4 is a locus with a rich repertoire of genetic variations, some of which are not discussed in this paper. In the future, it will be useful to measure and improve Kivvi’s accuracy in calling mosaic contracted alleles, in-cis duplications of D4Z4 arrays^39,40^ and D4Z4 proximal extended deletions (DPED alleles)^41^. In addition, we are extending Kivvi to genotype more clinically relevant VNTRs with large repeat units similar to D4Z4. One example that is already included in Kivvi is the Kringle IV-type 2 (KIV-2) repeat in LPA, where the KIV-2 copy number is associated with cardiovascular disease risks^42,43^. The development of more targeted informatics solutions for challenging genomic regions with HiFi data will continue to advance the consolidation of multiple genetic tests into a single workflow.

## Materials and methods

### Kivvi: HiFi-based tool for genotyping D4Z4

#### Read realignment

Kivvi accepts a genome aligned WGS BAM file (aligned to GRCh38) as input. It first extracts all reads overlapping D4Z4 on either chromosome 4 or chromosome 10 (chr4:190065229-190093263, chr4:190173122-190175903, chr10:133664430-133685491, chr10:133740609-133761980, GRCh38) and realigns them to a single D4Z4 repeat unit with Minimap2^44^. The reference for realignment is a 3298bp sequence extracted from CHM13-T2T reference genome with modifications, plus 905bp sequence distal to D4Z4 containing the polyA signal (Figure 1B).

#### Identification of unique repeat units and assembly into alleles

After realignment, Kivvi identifies unique copies of the D4Z4 repeat unit, which have distinct combinations of variants (Figure 1B). A de Bruijn graph is constructed where nodes represent distinct repeat units and edges represent links between distinct repeat units, extracted from HiFi reads, which typically contain 3 to 5 D4Z4 units per read. All D4Z4 alleles start with a partial unit, termed the starting unit, which can be identified as read segments softclipped at position 2058 (shorter read segments at the top right in Figure 1B). Read segments that align to D4Z4 and continue to the distal region containing the polyA signal indicate ends of qA alleles. qB alleles do not continue to the distal region containing the polyA signal. Instead, qB alleles end with a partial unit, which can be identified as read segments softclipped at position 429 (shorter read segments at the top left in Figure 1B). The nodes of the de Bruijn graph are assembled into alleles (Figure 1C). A complete D4Z4 allele refers to a path connecting the starting unit to the ending unit (either qB ends or the qA ends). A path that contains either the starting unit or the ending unit but not both is a partially assembled allele. For both fully assembled alleles and partially assembled alleles containing the ending unit, Kivvi reports the allele type (qAIntactPolyA, qB or qADisruptedPolyA) depending on the distal sequence and the status of the polyA signal. qAIntactPolyA and qADisruptedPolyA alleles both continue to the distal region. qAIntactPolyA alleles have intact polyA signal while qADisruptedPolyA alleles have mutations disrupting the polyA signal. Note that conventional 4qA alleles and the less common 4qA-L alleles^45,46^ are both reported as qAIntactPolyA alleles by Kivvi, as they both have the intact polyA signal. 4qA-L alleles can be identified by the 1.6kb insertion at the end of the last repeat unit (Figure S11).

#### Phasing of upstream haplotypes for chromosome 4/10 assignment

In order to distinguish chromosome 4 alleles from chromosome 10 alleles, Kivvi uses Paraphase^34^ to phase a 42kb genomic region that is homologous between chromosome 4 and chromosome 10 upstream of D4Z4. For this process, reads are extracted from chr4:190022510-190065229 and chr10:133622567-133664430 (GRCh38) and phasing is done for chr4:190022510-190065229. Four distinct haplotypes are identified for a sample, two from chromosome 4 and two from chromosome 10 (Figure 1D). The homology between chromosome 4 and chromosome 10 ends at chr4:190023009, and chromosome 10 haplotypes are softclipped at this position and thus shorter than chromosome 4 haplotypes. Kivvi identifies the two shorter haplotypes as chromosome 10 alleles and the two longer haplotypes as chromosome 4 alleles (Figure 1D). These haplotypes are connected to assembled D4Z4 alleles by checking overlapping reads so that Kivvi can report the originating chromosome of a D4Z4 allele.

#### Partially assembled D4Z4 alleles

If assembly of a complete D4Z4 allele fails due to the complexity of the repeat structure, Kivvi reports two partial alleles for each D4Z4 allele: 1) one partial allele (proximal) starts with the starting unit and connects with the upstream haplotypes, allowing assignment of the originating chromosome and 2) one partial allele (distal) that ends with the ending unit, allowing assignment of the allele type (qAIntactPolyA, qB or qADisruptedPolyA).

For each repeat unit on each partial allele, Kivvi assigns it to qADisruptedPolyA-like, qB-like or qAIntactPolyA-like units based on its variants, with the following requirements: qADisruptedPolyA-like (at least two of 256:C>T, 583:C>T and 683:A>G), qB-like (at least one of 172:A>G, 513:C>A and 2891:T>C) and qAIntactPolyA-like (none of those variants). The six variants are marked in Figure 1B. The variant signatures of the three different repeat unit types were extracted from the PCA in Figure 4A (also see Figure S5B and variant frequencies in Table S3).

After assignment of repeat units, Kivvi assigns each partial allele to one of the three allele types if more than 80% of its repeat units support the same allele type. After assignment of partial alleles, if there is a pair of proximal and distal partial alleles with the same assigned allele type in a sample (an unambiguous one-to-one mapping), these two partial alleles are joined into one allele and the sum of the sizes of both partial alleles is reported as the lower bound of the size of this merged allele. Kivvi reports the originating chromosome and allele type of this merged allele.

If a distal partial allele cannot be merged with a proximal partial allele, it is reported by Kivvi as a partial allele. The minimum of all starting allele sizes is added to its size and the sum is reported as the lower bound of array size for this partial allele. Kivvi reports the distal allele type of this partial allele, and the originating chromosome is reported as unknown as there is no information about the proximal end of this partial allele.

#### Methylation calling

Kivvi uses 101 CpG sites (Figure 1B), which are consistently highly methylated across samples, selected from within the D4Z4 repeat unit reference. In order to calculate the methylation value of a CpG site within a distinct repeat unit in a sample, Kivvi identifies all read segments that correspond to this repeat unit and calculates the median of the methylation values of those read segments at the CpG site. The methylation value of a repeat unit is calculated as the median methylation value of all 101 CpG sites within the repeat unit.

The reported methylation value of an allele is calculated as the median of all 101 CpG sites of the last five repeat units (or all repeat units if the allele is shorter than five units). As a sample-level summary metric, a median methylation value of all 101 CpG sites across all D4Z4 units is reported for each sample to represent the overall D4Z4 methylation level.

### Clinical samples and PacBio HiFi WGS sequencing

We collected 10 clinical samples with an FSHD1 diagnosis. The samples were shipped to GeneDx for analysis using the Bionano Saphyr system (Optical Genome Mapping). Ultra-high molecular weight DNA was extracted from either frozen cell pellet or blood (fresh or frozen) using the Bionano DNA extraction kit. The dsDNA was labeled with green fluorescent dye using Direct Label Enzyme, targeting specific sequence motifs with high sequence specificity and efficiency. In addition, the DNA backbone was labeled with blue dye. This allowed for imaging of sequence specific motifs on mega base-length gDNA by the Saphyr instrument. Results were analyzed using the Bionano EnFocus FSHD analysis tool according to the manufacturer’s protocol.

For HiFi WGS sequencing, genomic DNA was size-selected for long-read genome sequencing using PacBio SRE (catalog number 102-208-300), followed by shearing to a target size of 15-18 kb using the Hamilton Microlab STAR (Hamilton, Franklin, Massachusetts) system. Libraries were prepared with the PacBio SMRTbell® prep kit 3.0 (catalog number 102-141-700) and sequenced on one SMRT® Cell on a PacBio Revio® system with 24-hour movie times (targeted at generating 30x coverage).

HiFi WGS data was processed through the PacBio human WGS WDL pipeline (https://github.com/PacificBiosciences/HiFi-human-WGS-WDL). Specifically, alignment to GRCh38 was performed with pbmm2 v1.17.0 (https://github.com/PacificBiosciences/pbmm2), and small variants were called with DeepVariant^47^ v1.9.0.

In addition to the 10 FSHD1 samples, candidate pathogenic variants in *SMCHD1* were identified in 6 additional clinical samples (FSHD status unknown) using the above mentioned bioinformatics pipeline. These 6 samples were included as another dataset in this study.

### Population samples

We collected 601 unrelated population samples from five ancestral populations (291 Europeans, 95 Africans, 114 Mix Americans, 54 East Asians and 47 South Asians), collected from the Human Pangenome Reference Consortium (HPRC)^48,49^ and internal sample collections.

### Calculating two QC metrics for assessing data quality

Read length and read depth are the two key factors affecting the assembly of repeats and are thus used in the data quality comparison in Figure S1. The read length metric is calculated as the median of read lengths of all reads containing D4Z4 units. D4Z4 is a highly GC-rich repeat and its depth could be different from the genome average depth. The D4Z4 per-allele read depth is calculated as the number of reads overlapping the first partial repeat unit (Figure 1B) divided by four.

### Comparing Kivvi called alleles against orthogonal assays

Kivvi reports the size (exact for fully assembled alleles or lower bound for partial alleles), originating chromosome and distal allele type for each D4Z4 allele. There are four Kivvi reported alleles and four truth alleles by orthogonal assays (PFGE-Southern blot or OGM) in a sample, and they have to be mapped to each other before size comparison. A Kivvi allele is mapped to a truth allele with matching chromosome and allele type. When the reported chromosome is unknown for a Kivvi allele, qAIntactPolyA alleles are assigned to chromosome 4 truth alleles and qADisruptedPolyA alleles are assigned to chromosome 10 truth alleles. When more than one match with lower bounds is available, e.g. two chromosome 4 qAIntactPolyA alleles each in the truth data and Kivvi data (both partial alleles), truth alleles and Kivvi alleles are ordered by size in decreasing order and the longest truth allele is matched to the longest Kivvi allele, etc.

PFGE-Southern blot identified two haplotypes as 4qC, while they were reported as qAIntactPolyA by Kivvi. qAIntactPolyA and 4qC differ by just a few bases, none of which affect the polyA signal, so qC can be considered a subtype of qAIntactPolyA^35^.

Therefore, Kivvi does not report qC haplotypes, and these two haplotypes were considered correctly classified.

For allele size evaluation in Table 1, a size match is considered as a size difference between Kivvi and truth of three or less. A truth allele of 1-10RU is assigned “assembled with correct size” if the matching Kivvi allele is fully assembled with a matched size. It is considered inconclusive if it’s not fully assembled. A truth allele of >10RU is assigned “assembled with correct size” if the matching Kivvi allele is fully assembled with a matched size. A truth allele of >10RU and partially assembled by Kivvi is considered correctly classified if the reported lower bound is longer than 10, and is otherwise considered inconclusive.

### Mapping upstream sequences to known haplotypes

We selected major haplotypes from Figure 4 of Ref 35^35^ to use to label the upstream haplotypes in our population data. These haplotypes can be differentiated using bases at 10 genomic positions in the D4F104S1 region on chr4 (GRCh38): 190064812, 190064902, 190064922, 190064933, 190064937, 190064995, 190065046, 190065069, 190065077, and 190065157, together with genotypes of the (CA)_10_AA(CA)_11_ repeat in the SSLP region, which can be inferred from the sequence length of each haplotype between chr4:190061953-190061996. Bases at the 10 genomic positions were directly extracted from Kivvi output. The sequence length of the CA repeat was calculated by going through the Paraphase output BAM and counting the length of this region on each read supporting a haplotype, followed by taking the mode of this value for all reads of each haplotype. Finally, an upstream haplotype identified by Kivvi is assigned to a known haplotype by matching the sequence at the 10 positions and the length at the CA repeat as follows: 4A166/10A166: GCTGTGGAAG, 36bp; 4A168/4B168: GCTGTGGAAG, 38bp; 4A161: CTCACATTGT, 42bp; 4A159: CTCACATTGT, 40bp; 4B163: CTTACATTGT, 44bp; 10B161: CTTACATTGT, 42bp; 10A176: GCTGTGGAAG, 46bp; 4B162: GCTGTGGAAG, 32bp.

### Separating upstream haplotypes into four groups

In order to define simpler haplotype groups for the upstream region, we performed PCA (Figure S8) of upstream haplotypes in European, East Asian and South Asian populations to exclude hybrid alleles. Three chromosome 4 alleles ending in qADisruptedPolyA were removed prior to PCA. Upstream haplotypes were separated into two major groups on the PC1 axis (Figure S8, left: Group2; right: Group1). Base positions that are always different between Group1 and Group 2 were selected to identify the two major groups. Group1 is further divided into two subgroups as separated on the PC3 axis in Figure S8C (upper: Group1.1, lower: Group1.2). Group2 is further divided into two subgroups as separated on the PC2 axis in Figure S8B (upper: Group2.1, lower: Group2.2). Within each group, base positions that are always different between subgroups were selected to identify the subgroups. In total, ten base positions (chr4, GRCh38: 190064271, 190064812, 190064902, 190064922, 190064933, 190064937, 190064995, 190065046, 190065069, and 190065077) were selected to identify the four groups as follows: Group1.1: TCTCACATTG; Group1.2: TCTTACATTG; Group2.1: TGCTGTGGAA; Group2.2: AGCTGTGGAA.

### Principal component analysis

Principal component analysis (PCA) was conducted on single nucleotide variant (SNV) sites identified across all sequences using the prcomp function in R.

## Data and code availability

HiFi WGS data for HPRC samples are documented in https://github.com/orgs/human-pangenomics/repositories and can be downloaded from https://s3-us-west-2.amazonaws.com/human-pangenomics/index.html?prefix=working/ for analysis.

Kivvi is available on https://github.com/PacificBiosciences/kivvi. Kivvi Version 1.0 was used in all analyses of this paper.

## Declaration of interests

XC, ZK, ED and MAE are employees of Pacific Biosciences. JMD, JN, ASB, SY, SL, KN and ASL are employees of GeneDx. SMvdM is a board member of Renogenyx and has acted as consultant and/or is member of the advisory board for several pharmaceutical companies developing therapeutics for FSHD. SMvdM and RJLFL are coinventors of several FSHD-related patent applications.

## Acknowledgments

We thank the Human Pangenome Reference Center (HPRC) for generating and releasing the HiFi WGS data. We thank Billy Rowell for help with data transfer and storage.

## Notes

https://github.com/PacificBiosciences/kivvi

