## Supplementary figures for "Resolution of the D4Z4 repeat responsible for facioscapulohumeral muscular dystrophy with HiFi sequencing"

1. PacBio, Menlo Park, CA, USA.
2. Department of Human Genetics, Leiden University Medical Center, 2333 ZA Leiden, Netherlands.
3. GeneDx, Gaithersburg, MD, USA.

### Supplementary Figures

Figure S1. Median read length and mean coverage per allele in D4Z4 for 1kGP samples, FSHD1 patient samples and samples with *SMCHD1* pathogenic variants.

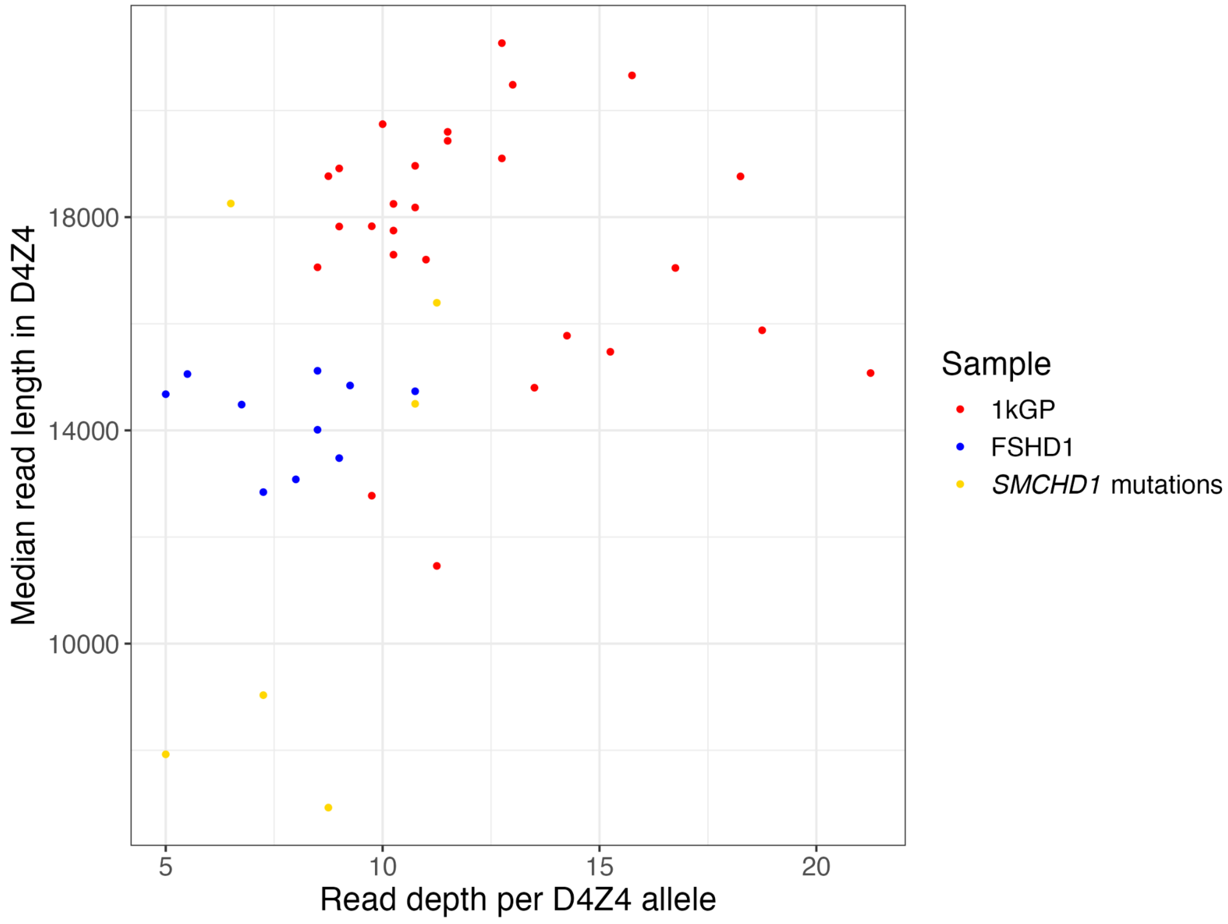

Figure S2. Pairwise distance between D4Z4 repeat units, comparing a pair of repeat units in the same population (top), on the same allele (middle) and next to each other on the same allele (bottom).

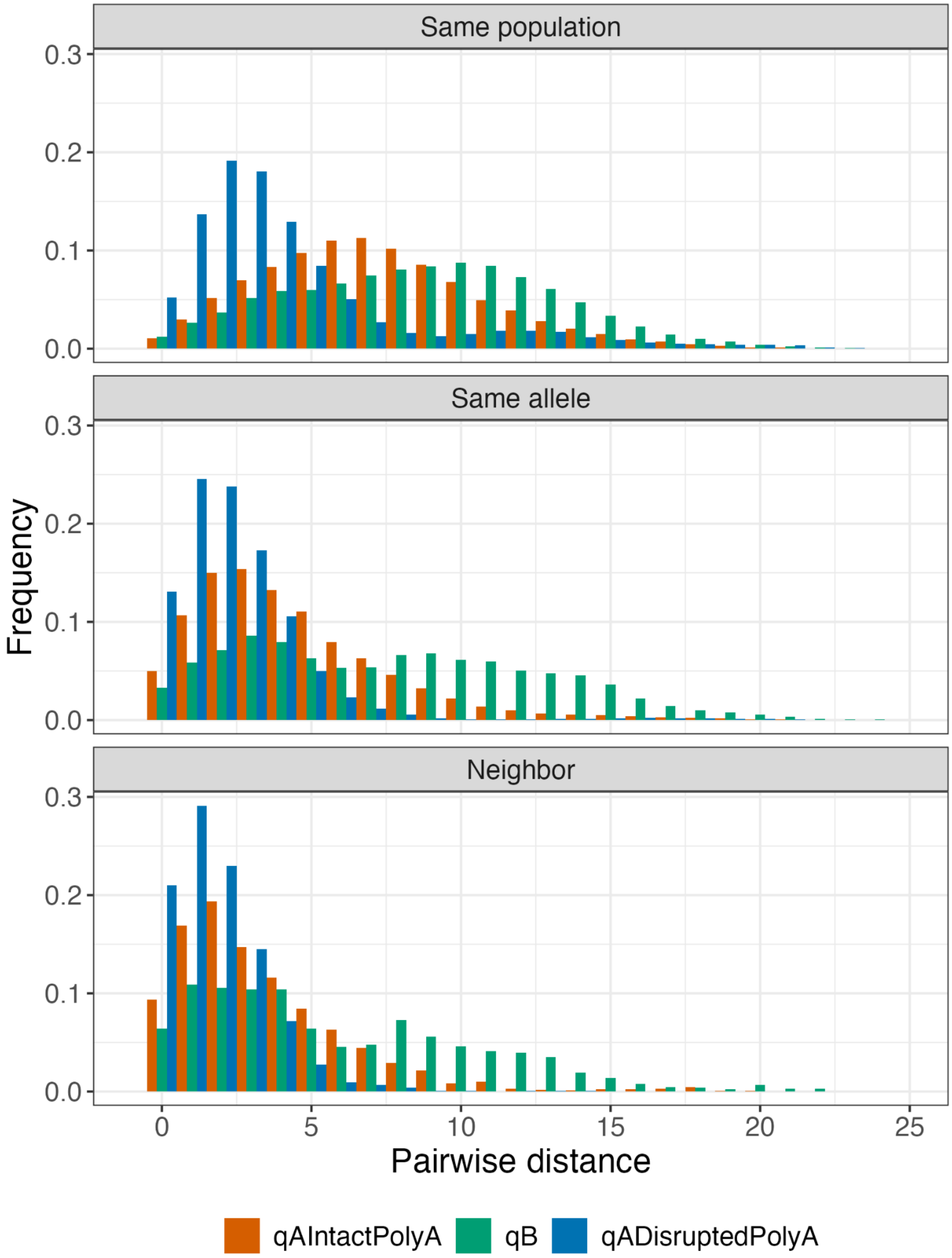

Figure S3. Distribution of qAIntactPolyA, qB and qADisruptedPolyA alleles on chr4 and chr10 among fully assembled D4Z4 alleles.

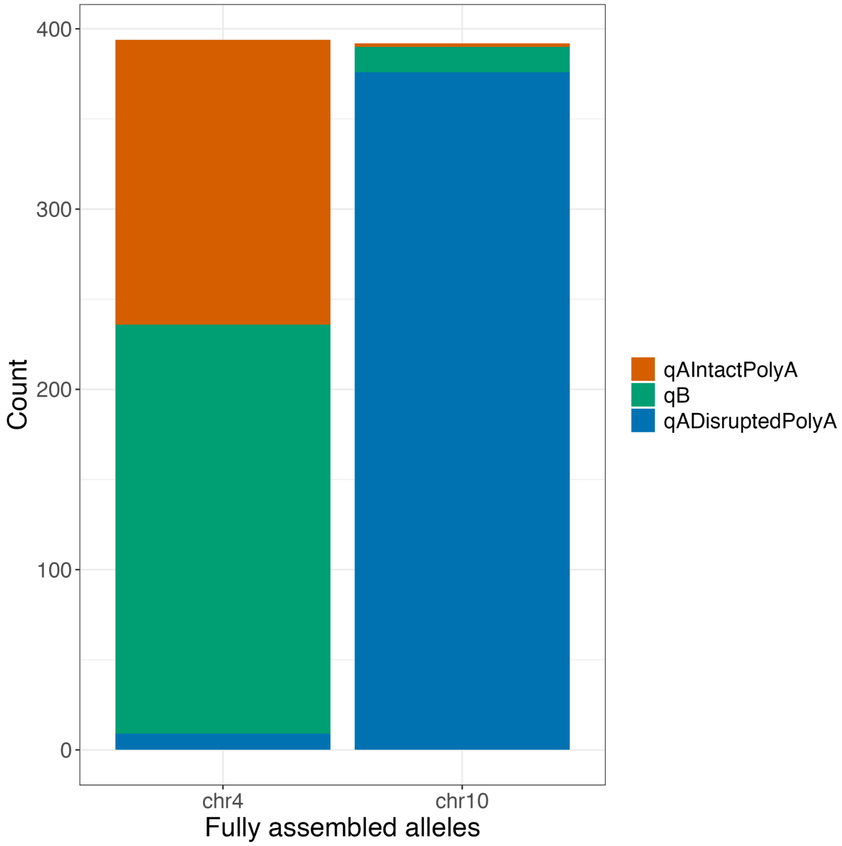

### Figure S4. *SMCHD1* pathogenic variants identified in codon V615 (first codon after splice site) in sample GeneDx016 and GeneDx001, marked by black arrows.

This codon is right next to the splicing site and thus the two variants may potentially affect splicing. The A>G variant nearby is a known polymorphism (rs635132).

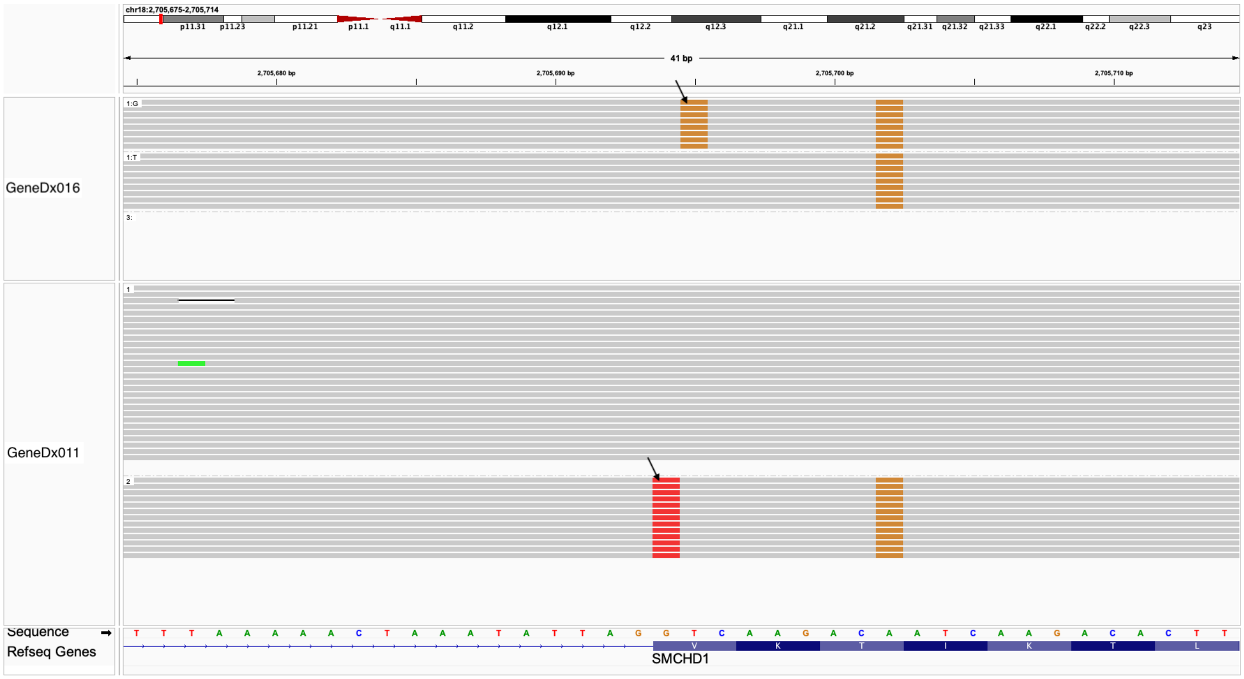

### Figure S5. Details for the PCA of D4Z4 repeat units in Figure 4A.

**A.** Plot of percent variation explained. **B.** Same PC1 vs. PC2 plot as in Figure 4A, with boxes outlined in gray indicating different types of repeat units based on variant profiles: (from left to right) qADisruptedPolyA-like, qAIntactPolyA-qADisruptedPolyA hybrid, qAIntactPolyA-like, and qB-like.

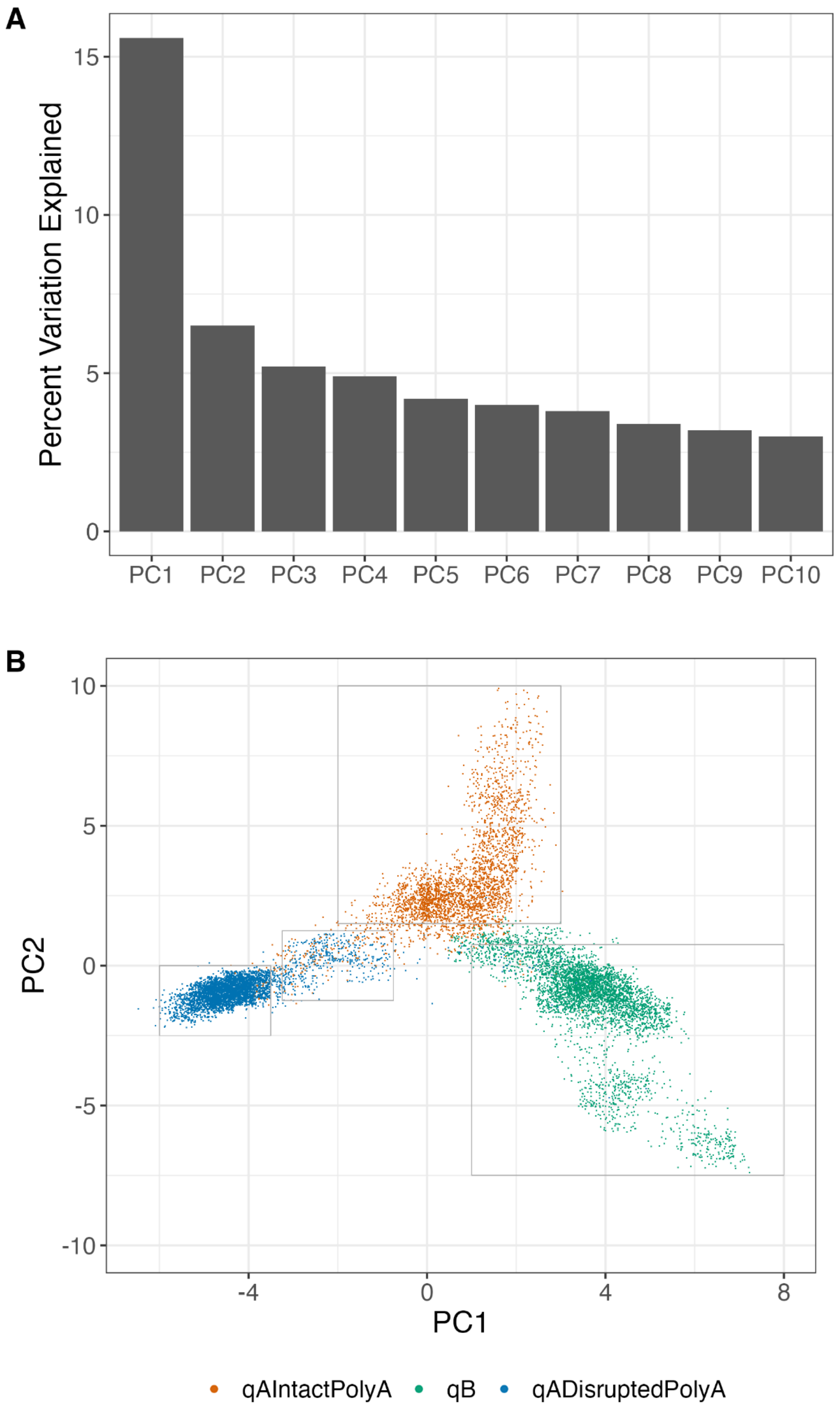

### Figure S6. PCA of D4Z4 repeat units (same as in Figure 4A and Figure S5B) grouped by chromosome and the status of the BlnI (B) and XapI (X) restriction sites.

The gray-bordered box denotes the qAIntactPolyA-qADisruptedPolyA hybrid zone.

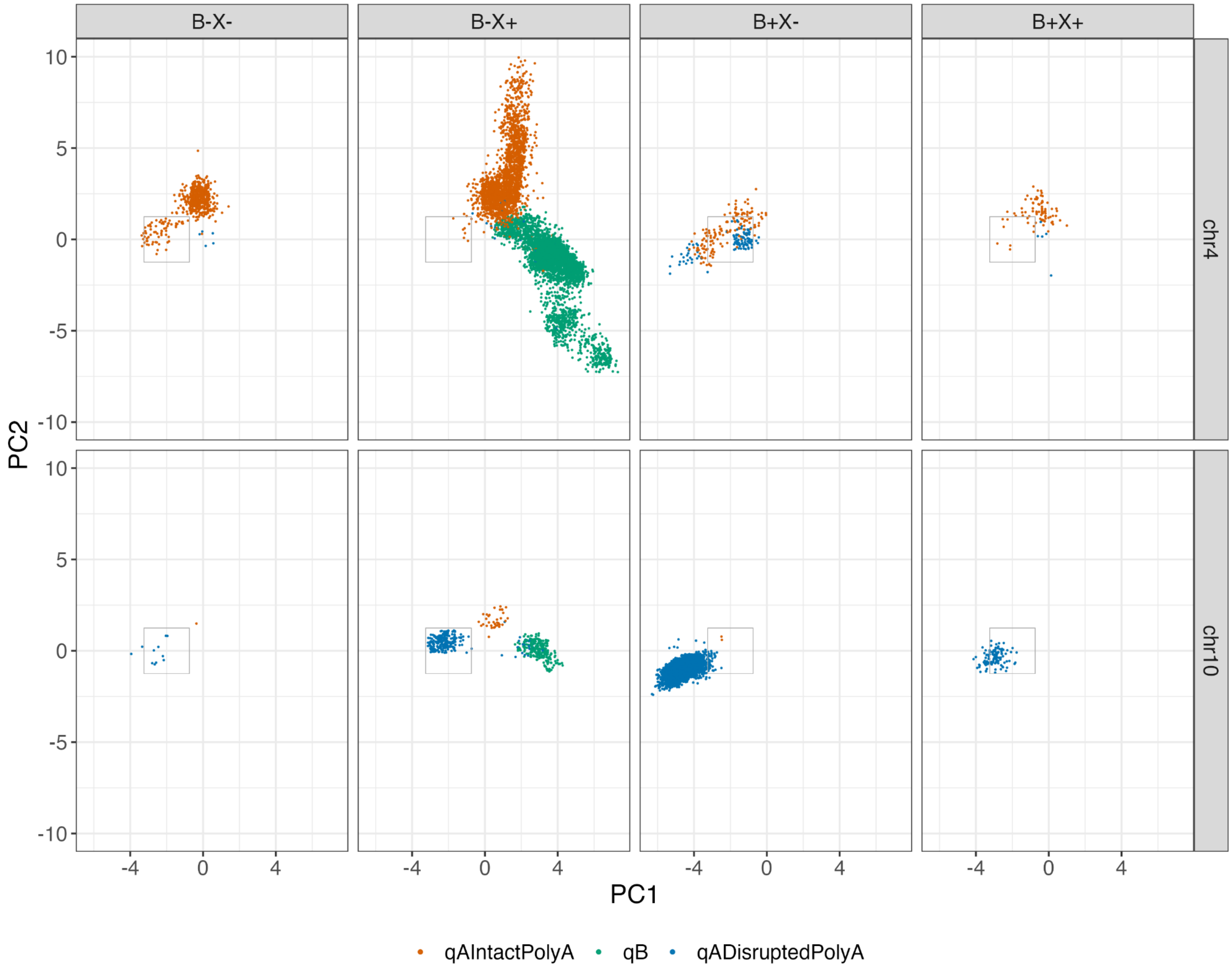

Figure S7. PCA of haplotypes from a 3640bp region upstream of D4Z4.

**A.** Plot of percent variations explained. **B.** Same PC1 vs. PC2 plot as in Figure 4B. Colors represent previously identified haplotypes as labeled in panel C. **C.** Same PC1-PC2 plot as B, grouped by distal allele types and originating chromosomes.

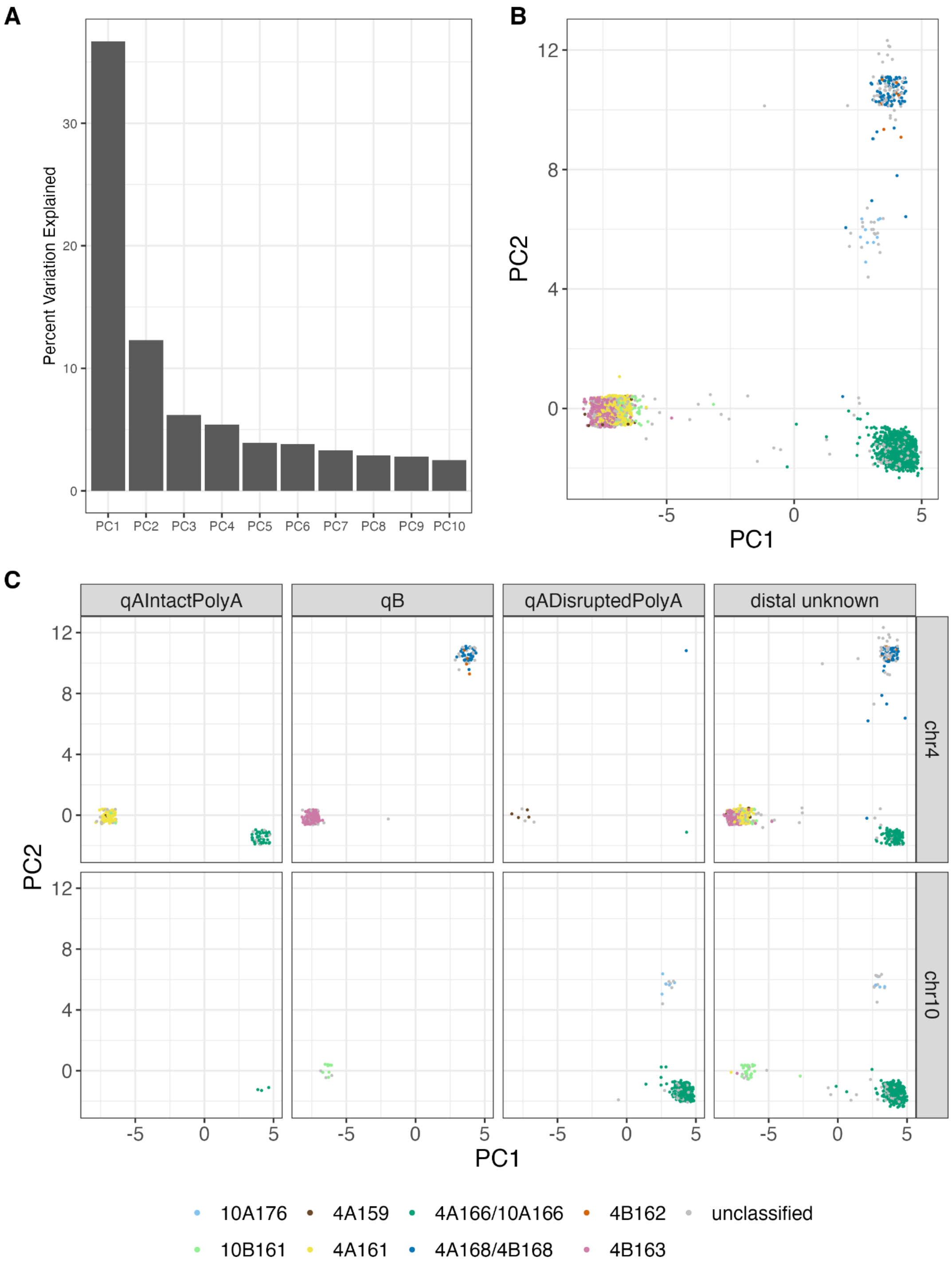

### Figure S8. PCA of upstream haplotypes reveals separation of Group1 and Group2 on PC1, separation of qA and qB in Group2 on PC2 and separation of qA and qB in Group1 on PC3.

PCA of upstream haplotypes in EUR, EAS and SAS populations only (after removal of three chr4-qADisruptedPolyA alleles). **A.** Plot of percent variation explained. **B.** PC1 vs. PC2. Left is Group2 and right is Group1. Within Group2 (left), upper is Group 2.1 and lower is Group 2.2. **C.** PC1 vs. PC3. Within Group1 (right), upper is Group1.1 and lower is Group1.2.

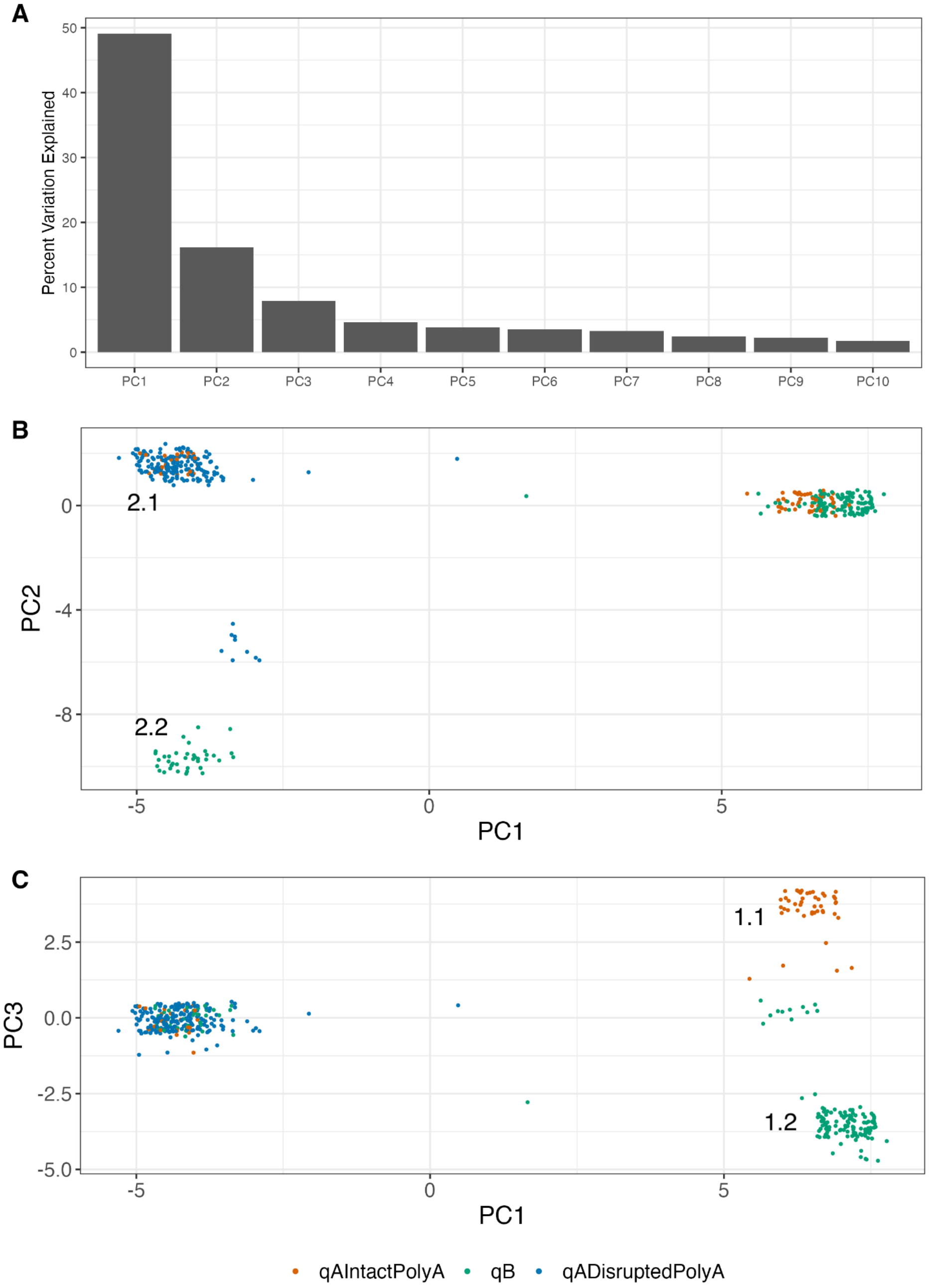

### Figure S9. Translocation alleles show transitions in repeat unit signatures indicating breakpoint.

Each subplot shows the repeat units of one translocation allele. Units from the most proximal to the most distal are represented using the lightest to darkest colors. On chr4 Group1.2 (left) and chr4 Group2.2 (right), qB alleles are expected. Both alleles are a qADisruptedPolyA allele that starts with qB-like units and transitions to qADisruptedPolyA-like units. The transition point indicates the translocation breakpoint. The gray-bordered box denotes the qAIntactPolyA-qADisruptedPolyA hybrid zone.

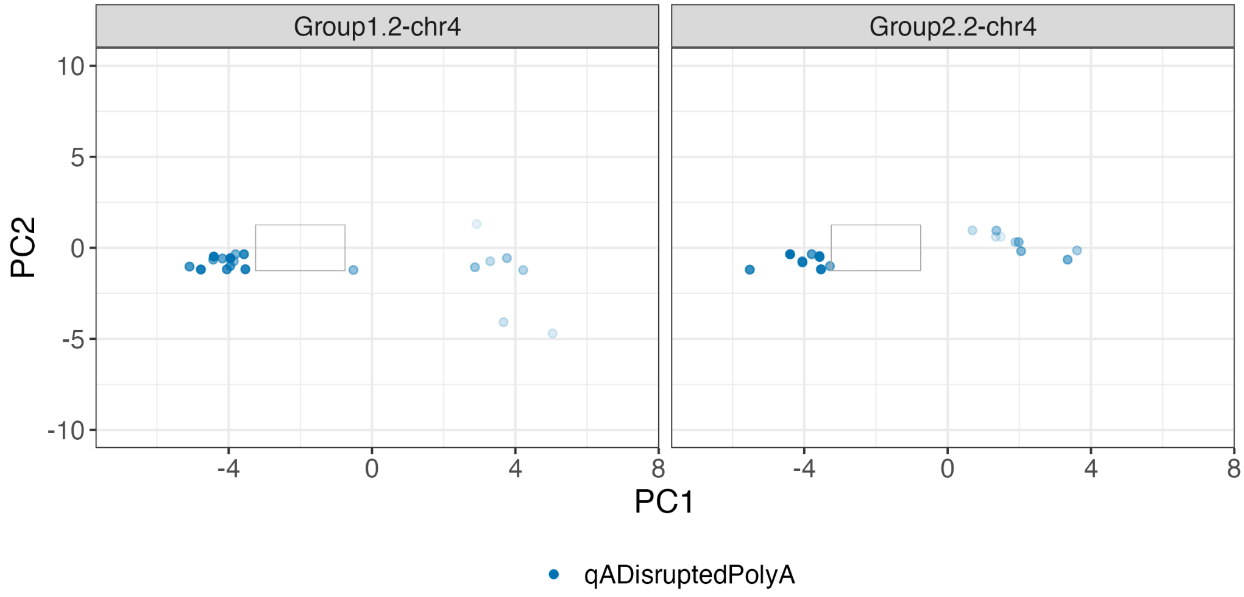

### Figure S10. qADisruptedPolyA alleles on chr10 Group2.2 often show qAIntactPolyA-qADisruptedPolyA hybrid units at the proximal end and qB-like units at the distal end.

Each subplot shows the repeat units of one allele. Units from the most proximal to the most distal are represented using the lightest to darkest colors. The gray-bordered box denotes the qAIntactPolyA-qADisruptedPolyA hybrid zone.

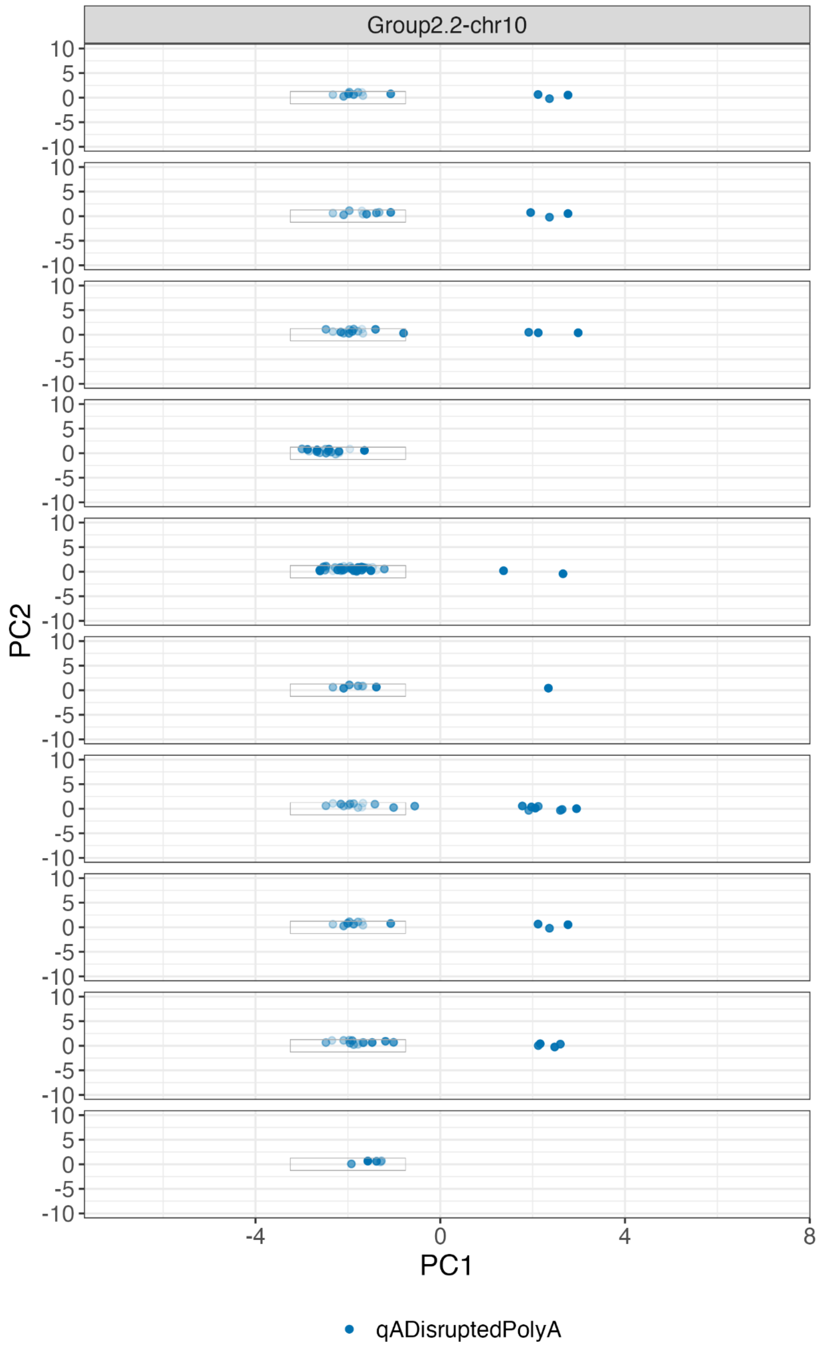

Figure S11. 4qA-L alleles show a 1.6kb insertion at the end of the last repeat unit (RU index 7 in sample NA19650).

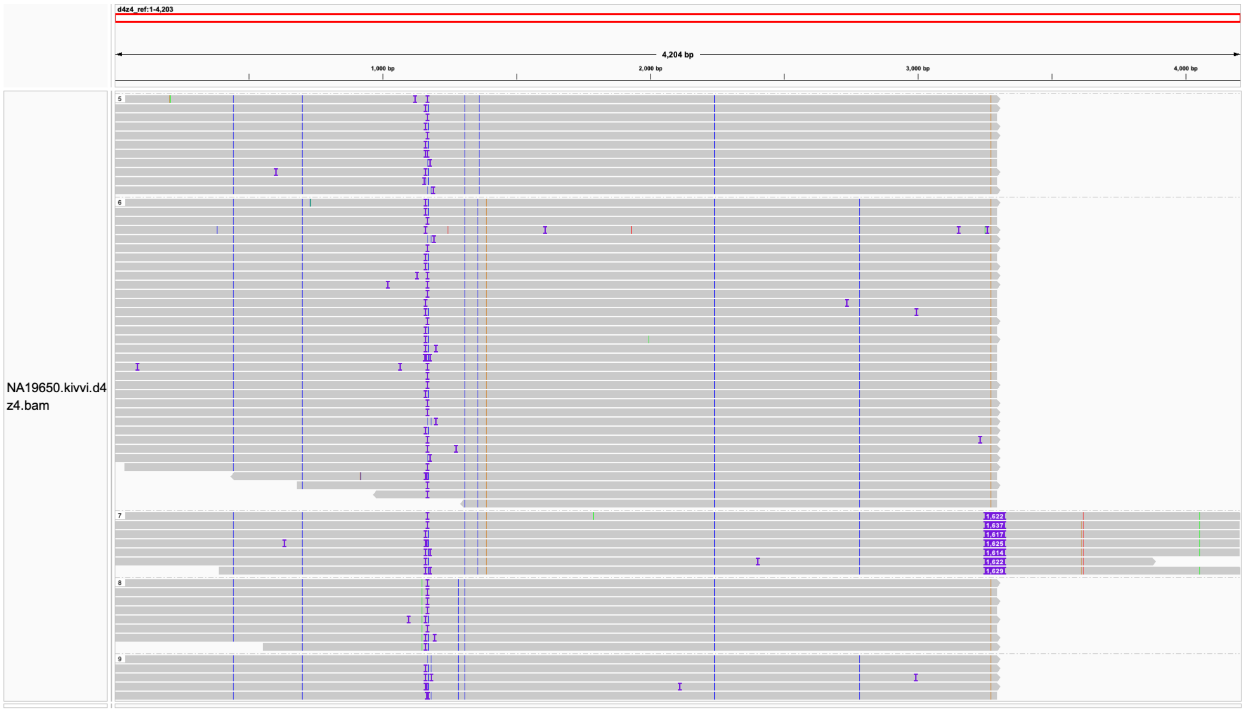

### Supplementary Tables

Table S1. Details of D4Z4 calls in 27 1kGP samples with PFGE-Southern blot results. (Excel Spreadsheet)

Table S2. Details of D4Z4 calls in 10 FSHD1 patient samples. (Excel Spreadsheet)

Table S3. Variant frequencies in different types of repeat units (defined by the PCA in Figure S5B).
